# Scalable environmental DNA methods reveal strong associations between landscape-scale forest habitat and insect richness

**DOI:** 10.1101/2025.04.01.645213

**Authors:** Rodney T. Richardson, Grace Avalos, Cameron Garland, Regina Trott, Olivia Hager, Mark J. Hepner, Clayton Raines, Karen Goodell

**Affiliations:** Western EcoSystems Technology, Inc., Indianapolis, IN; Department of Entomology, University of Minnesota, Saint Paul, MN; Appalachian Laboratory, University of Maryland Center for Environmental Science, Frostburg, MD; Metamorphic Ecological Research and Consulting, LLC, Alonzaville, VA; U.S. Geological Survey, Eastern Ecological Science Center, Leetown, WV; Department of Evolution, Ecology and Organismal Biology, The Ohio State University, Newark, OH

**Keywords:** Invasion biology, Entomology, Conservation, Ecology, Biodiversity

## Abstract

1. While aquatic environmental DNA (eDNA) methods have reached relative maturity, terrestrial eDNA methods are nascent and have yet to reach widespread use. Field-ready applications require eDNA survey methods where samples are easy to collect by inexperienced practitioners, easy to transport between the field and lab, and easy to process thereafter. Here, we demonstrate methods that satisfy these requirements and show strong potential for characterizing diverse terrestrial eDNA samples collected from flower and leaf surfaces.
2. We used novel methods to collect and process 236 flower eDNA samples and 21 leaf surface eDNA samples, obtaining 2,228 Arthropoda eDNA detections spanning 175 families using amplicon sequencing of two genetic markers.
3. Detected taxa were diverse and included numerous groups of conservation concern, such as bees (Hymenoptera; Anthophila, 32 genera spanning 5 families) and Lepidoptera (209 genera from 21 families). Data reveal strong associations between insect community richness and remotely sensed measures of forest habitat, providing a quantitative perspective of relevance to insect conservation.
4. It is increasingly clear that a variety of organisms readily disperse eDNA throughout the environment, supporting the notion that eDNA will be a powerful tool for characterizing species distributions and monitoring at-risk species. However, we conclude that researchers seeking to characterize fine-scale habitat associations or plant-pollinator interactions using eDNA will need to carefully design studies with appropriate field controls, such as the leaf surface eDNA samples collected here.

## Introduction

In recent decades, North America has experienced considerable shifts in land use, with more than 100,000 km^2^ converted into urban developments and row crop agriculture (He et al., 2019; Lark et al., 2020). Areas occupied by short vegetation, including shrublands and grasslands, have decreased by over 200,000 km^2^ (Song et al., 2018). Land use changes, and other forms of global environmental change, are intrinsically linked to change in the abundance and geographic distribution of species (Jackson & Sax, 2010). Such changes represent a direct risk to the ecosystem processes that generate natural capital humans depend upon. Accordingly, scientists have argued for rapid investment in the infrastructure and methods needed to better understand the effects of environmental change as well as how they can be managed to limit economic damage and improve ecosystem resilience (Williams, 2022). Attaining these goals requires methods to track shifts in abundance, range and diversity of key biota that underly ecosystem function.

One of the few encouraging attributes of recent land use change includes the reforestation of over 270,000 km^2^ across the U.S. and Canada (Song et al., 2018). Forests can help buffer against extreme conditions associated with climate change (Barnes et al., 2024; De Lombaerde et al., 2022) and provide valuable habitat for a wide diversity of wildlife, especially when differential management is used to promote variation in forest age, composition and understory conditions (Hekkala et al., 2023; Hepner et al., 2024; Jeffries et al., 2006; Uhl et al., 2024). Insect community trends in forested systems are difficult to measure, but the maintenance of forest appears to be linked to lower magnitudes of insect loss (Martínez-Núñez et al., 2024; Staab et al., 2023) and, potentially, diversity gains (Ober & Hayes, 2008).

Insects exert major influence over terrestrial ecosystems and are sensitive to environmental change (Jactel et al., 2019). Insects are of broad conservation concern because recent reports suggest insect biomass and richness are declining (Hallmann et al., 2017; Sánchez-Bayo & Wyckhuys, 2019; Wagner et al., 2021), but efforts to characterize such declines have come to different conclusions about the strength of the evidence and rates of loss (Crossley et al., 2020; Fürst et al., 2023; Rosenheim & Ward, 2020). Meanwhile, there is mounting concern regarding climate change-associated insect pest outbreaks threatening forest health (Bracalini et al., 2024). Disagreement over the magnitude and associated drivers of insect population changes indicates that studies with greater statistical power and improved design are needed. Insects, however, are difficult to monitor because of their high diversity, small size and cryptic variation (van Klink et al., 2022). Even large datasets can fall short of the power needed to link insect compositional change with environmental factors (Tronstad et al., 2022), and the lethal nature of typical designs could compromise vulnerable populations (Tepedino et al., 2015).

In addition to the sheer number of samples needed for large-scale insect monitoring, variation in the properties of different collection methods limits comprehensiveness and comparability across studies (Didham et al., 2020; Montgomery et al., 2021; Portman et al., 2020). Thus, researchers often rely on multiple survey methods to minimize sampling bias, though the extent to which this increases inferential accuracy has received minimal attention. Once samples are collected using traditional methods, the preservation of individual specimens is cumbersome and subsequent identification with taxonomic keys requires high degrees of expertise (Packer et al., 2009). The difficulty of measuring insect population change hampers our ability to stay abreast of important shifts of relevance to insect conservation and forest health (Didham et al., 2020; Knuff et al., 2019; Kuhlman et al., 2021).

Environmental DNA (eDNA) methods have been proposed as a means of circumventing key challenges in the surveillance of terrestrial diversity (Allen et al., 2023; Bell et al., 2024; Bohmann & Lynggaard, 2023). Rapid collection of large numbers of samples can be accomplished by individuals with minimal training using eDNA sampling. These features allow surveys to be distributed across broad communities of scientists. Additionally, samples can be collected easily over short time periods to closely align surveys with the natural life history of target taxa. In this study, we demonstrate the advantages of eDNA sampling for characterizing insect diversity across a broad spatial scale in a heavily forested ecosystem that, like most forests globally, is experiencing the dual threats of climate and land use change. In sequencing collected eDNA samples using genetic markers with previously demonstrated sensitivity across a variety of insect orders (Avalos et al., 2024), we hypothesized that insect diversity would be positively related to the proportion of forested area and forest tree species diversity at the landscape scale. In addition to evaluating relationships between forest habitat and insect diversity, we provide a rough comparison of the sensitivity of our methods to past insect eDNA efforts.

## Methods

### Study area and Sample Collection

We conducted a large-scale eDNA survey of the Central Appalachian region of the Eastern United States (U.S.). Central Appalachia is a montane region with large areas of temperate forest and high forest connectivity (Belote et al., 2016). Unforested areas in this region tend to include pasture, row-crop agriculture and low-intensity development (Hepner et al., 2024). From June to September 2022, we conducted 71 site visits across 55 locations spanning 4 states in the Eastern U.S.: Pennsylvania, Maryland, West Virginia and Virginia (Figure 1A). Sites consisted of roadside rights-of-way with abundant floral resources. During each site visit, 1 to 3 surveyors collected flower and leaf surface eDNA samples. Leaf surface samples included leaves from trees or shrubs that were at least 3 meters from prominent floral forage, and which showed no evidence of insect herbivory or visible debris. For each eDNA sample, a single species of flower or leaves were carefully clipped into a sterile quart sized zip-closable plastic bag until the bag was approximately ¼ to ¾ full of loose plant material. Samples were mixed with 25 mL of eDNA preservative (10 percent EtOH v/v, 40 percent propylene glycol v/v and 0.25 percent sodium dodecyl sulfate w/v), mixed gently and allowed to sit for 1 to 5 min before the eDNA-containing preservative rinse was carefully poured into a labelled 50 mL conical vial for transport to the laboratory (Figure 1B). During eDNA collection, researchers used flame-sterilized tools and disposable sterile nitrile gloves to avoid cross contamination between sites or eDNA surveys.

**Figure 1:**
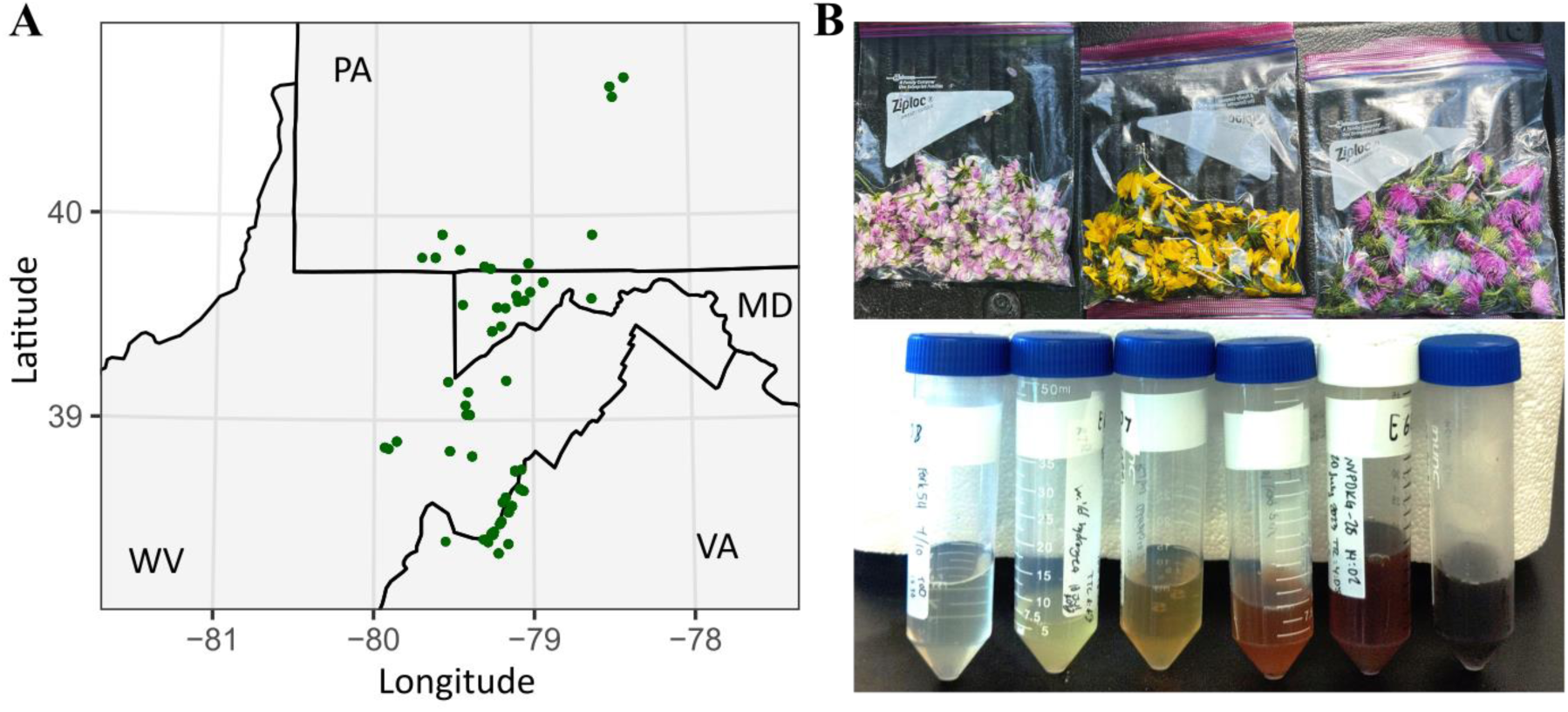
Map of eDNA sampling locations (A), and photos illustrating the sample collection and preservation methods used (B). Mapping was conducted using ggplot2 (Wickham, 2016) and photos used with permission from Andrew Held, Frostburg State University.

### Laboratory eDNA Processing

To purify each field-collected eDNA sample, 10 ul of 11.25 ug/ul yeast tRNA, 15 ul of 0.5 percent w/v linear polyacrylamide (Maixner et al., 2022) and 20 ul of 20 mg/ml glycogen was added to each sample. The concentration of NaCl was adjusted to 0.2 M through addition of 5 M NaCl stock solution. Samples were incubated overnight at −20° C to facilitate aggregation of nucleic acids prior to centrifugation at 7,006 x g for 40 min at 4° C. Sample supernatant was discarded and the eDNA-containing pellet was air-dried until all visible liquid was evaporated. Pellets were resuspended in 500 ul of lysis buffer (0.5 percent SDS, 10 mM Tris, 1 mM EDTA, 10 mM NaCl, pH 8.0). Resuspended samples were transferred to 2.0 mL microcentrifuge tubes and 5 ul of 20 mg/ml proteinase K was added prior to incubation at 55° C for 1 hour. Each sample was brought to a NaCl concentration of 0.3 M through addition of 5 M stock solution. Samples were incubated on ice for 5 min prior to centrifugation at 20,000 x g for 20 min at 4° C. The eDNA-containing supernatant was transferred to a new 2.0 ml microcentrifuge tube and approximately 2.5 volumes of ethanol were added. Samples were gently mixed by inversion and incubated at −20° C overnight prior to centrifugation at 20,000 x g for 40 min at 4° C. The supernatant was carefully removed, and the eDNA-containing pellets were air dried for approximately 15 minutes prior to being resuspended in 100 ul of nuclease-free water. PCR inhibitors were then removed from the DNA using the Qiagen PowerClean Pro Cleanup Kit. Following purification, Illumina MiSeq amplicon libraries were prepared using a previously-established 3-step PCR-based protocol (Richardson et al., 2019). During library preparation, we used two sets of primers, the 28S primers from Darby et al. (2020), designed for specificity to Anthophila, and the broad arthropod-specific BR2/BF2 primers (Elbrecht & Leese, 2017).

During initial BR2/BF2 PCR reactions, 1.6 uL of a 100 uM concentration equimolar mixture of blocking primers was added. These 3’ spacer C3 modified primers were designed to limit fungal byproducts previously observed in past work (Avalos et al., 2024) and are provided in supplemental Table S1. For each marker, we incorporated 15 no-library negative control dual-index combinations during library preparation to quantify critical-mistagging rates during downstream sequencing (Esling et al., 2015). Amplified libraries were purified with the SequalPrep Normalization Plate Kit, pooled equimolarly and subjected to Illumina MiSeq sequencing using a single-end 300 cycle v2 flow cell for 28S and paired-end 500 cycle v2 flow cell for COI.

### Taxonomic Annotation of eDNA Data

After sequencing, VSEARCH (Rognes et al., 2016) was used to merge paired-end COI sequences, quality trim single-end 28S sequences and remove priming sites from the 5’ and 3’ ends. During quality trimming, sequences were truncated starting with the first base pair with a Phred score of 18 or lower. Sequences were taxonomically annotated by semi-global VSEARCH top-hit alignment against custom reference sequence databases trimmed to the amplicon regions of interest using MetaCurator (Richardson et al., 2020), with dependencies HMMER (Eddy, 2011), MAFFT (Katoh et al., 2002), VSEARCH and Taxonomizr (Sherrill-Mix, 2019). During alignment, a minimum query cover of 0.8 was required. Reference databases were constructed using sequencies available through NCBI, downloaded on February 5^th^, 2023 for 28S and February 2^nd^, 2023 for COI. Following reference curation, the 28S database consisted of 53,478 sequences representing 1,289 families, 11,440 genera and 28,191 species. COI reference data were relatively more complete, with 606,748 sequences representing 2,129 families, 28,975 genera, and 104,243 species. During alignment, 96 percent identity matches were considered confident genus-level detections for 28S and 95 percent matches were considered genus-level detections for COI. For both markers, 85 percent matches were considered phylum-level matches. Since all Illumina sequencing runs produce low frequency misidentifications when inferring dual-index tags (i.e. critical mistags; Esling et al., 2015), we used an established statistical technique to remove identifications with greater than 0.05 probability of representing a critical mistag-associated detection (Richardson, 2022). Additionally, we removed detections represented by three or fewer sequences in a sample. Computational analysis was performed using the *Ohio Supercomputer Center*, (1987).

### Survey Site Landscape Analysis

For each sampling site, remotely sensed maps of land cover and forest species basal areas were used to gather landscape-scale information at 0.5, 1 and 3 km radius scales. Proportional land cover was characterized using the National Land Cover Dataset (NLCD; Dewitz, 2021) and downloaded for our study region using the get_nlcd function in the FedData package (Bocinsky, 2024) on February 5^th^, 2024. Total forest basal area and species-specific forest basal areas were estimated using a 250 m resolution gridded product based on USFS Forest Inventory and Analysis (FIA) data, MODIS remote sensing of phenology and environmental parameters (Wilson et al., 2012). From these data, we defined three predictor variables for modelling. *DeciduousForest* was represented by the NLCD Deciduous Forest category and *EvergreenForest* was calculated as the sum of the NLCD Evergreen Forest and Mixed Evergreen Forest categories. *ForestDiversity* was estimated by calculating Shannon diversity values (Shannon, 1948) from FIA-inferred species-specific basal areas.

### Statistical Analysis

Statistical analysis was conducted using R v4.3.2 (R Core Team, 2021). Following taxonomic sequence annotation, we aggregated all detections across the two genetic markers, producing a single table of presence-absence values. Leaf surface eDNA samples were then compared against flower eDNA samples using a Chi-square test to evaluate differences in frequency of eDNA detection per sample and a Wilcoxon Rank Sum test to evaluate differences in genus-level eDNA richness per sample. These tests were conducted for all Insects, as well as Anthophila and Lepidoptera, two taxa enriched for species of conservation concern. Since bee richness and frequency of detection was expected to be greater in flower eDNA relative to leaf surface eDNA samples, one-tailed tests were used when conducting analyses for Anthophila.

To evaluate landscape-scale diversity trends, we analyzed eDNA-derived richness estimates as a function of local forest cover and forest tree species diversity. For these analyses, a starting generalized linear mixed model was constructed by regressing eDNA richness per sample against the proportion of *DeciduousForest*, *EvergreenForest* and *ForestDiversity*. Terms for sampling week (*W*) to account for systematic trends in insect diversity over the sampling period and mean proportion of Arthropoda eDNA sequences (*D*), weighted by the total study-wide detections for each genetic marker to account for how intensively the sample was interrogated during sequencing, were included. To account for repeated measures, a random intercept term for substrate was added (Equation 1). For flower eDNA samples, substrate labels indicated the plant family from which the sample originated, while leaf surface samples were given a single unique factor label. *EvergreenForest* (3-km scale) values were z-scaled to circumvent model convergence issues. A Poisson error distribution was specified, and reverse stepwise selection was conducted where the least significant of all non-significant (α = 0.05) features was removed during each modelling round. Feature selection was first conducted with landscape predictors calculated at the 3-km scale, using the Watanabe-Akaike information criterion (WAIC; Watanabe, 2013) to select the best-supported model. We then tested the final model using predictors calculated at the 1- and 0.5-km scales. Following selection, each model was tested for overdispersion. Where necessary, confidence values were adjusted using recommendations associated with the lme4 package (Bates et al., 2015). Modelling was conducted for Arthropoda, Hymenoptera and Lepidoptera, all of which exhibited >400 genus-level detections.

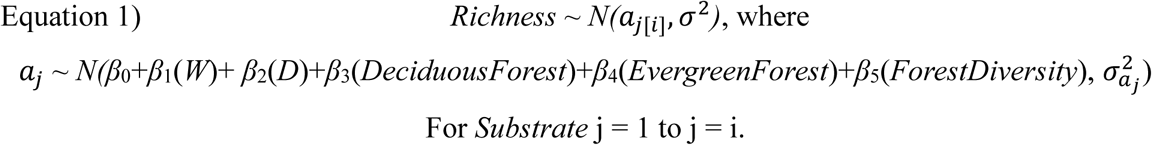

## Results

During field sampling, 236 flower and 21 leaf surface eDNA samples were collected. Following sequencing, 28S and COI libraries were sequenced to mean depths of 38,280 and 32,329 reads per sample, respectively (Figure 2A). The 28S marker showed moderate specificity for Arthropoda amplification, with a median of 10.9 percent of sequences per sample belonging to Arthropoda. However, COI exhibited considerable non-target amplification and a median of only 0.4 percent of sequences per sample represented Arthropoda (Figure 2B), suggesting that the blocking primers employed during amplification were not effective in improving reaction specificity. Similarly, while a median of 4.79 percent of total 28S sequences could be identified to an arthropod genus, only 0.08 percent of COI sequences were identifiable to this level (Figure 2C).

**Figure 2:**
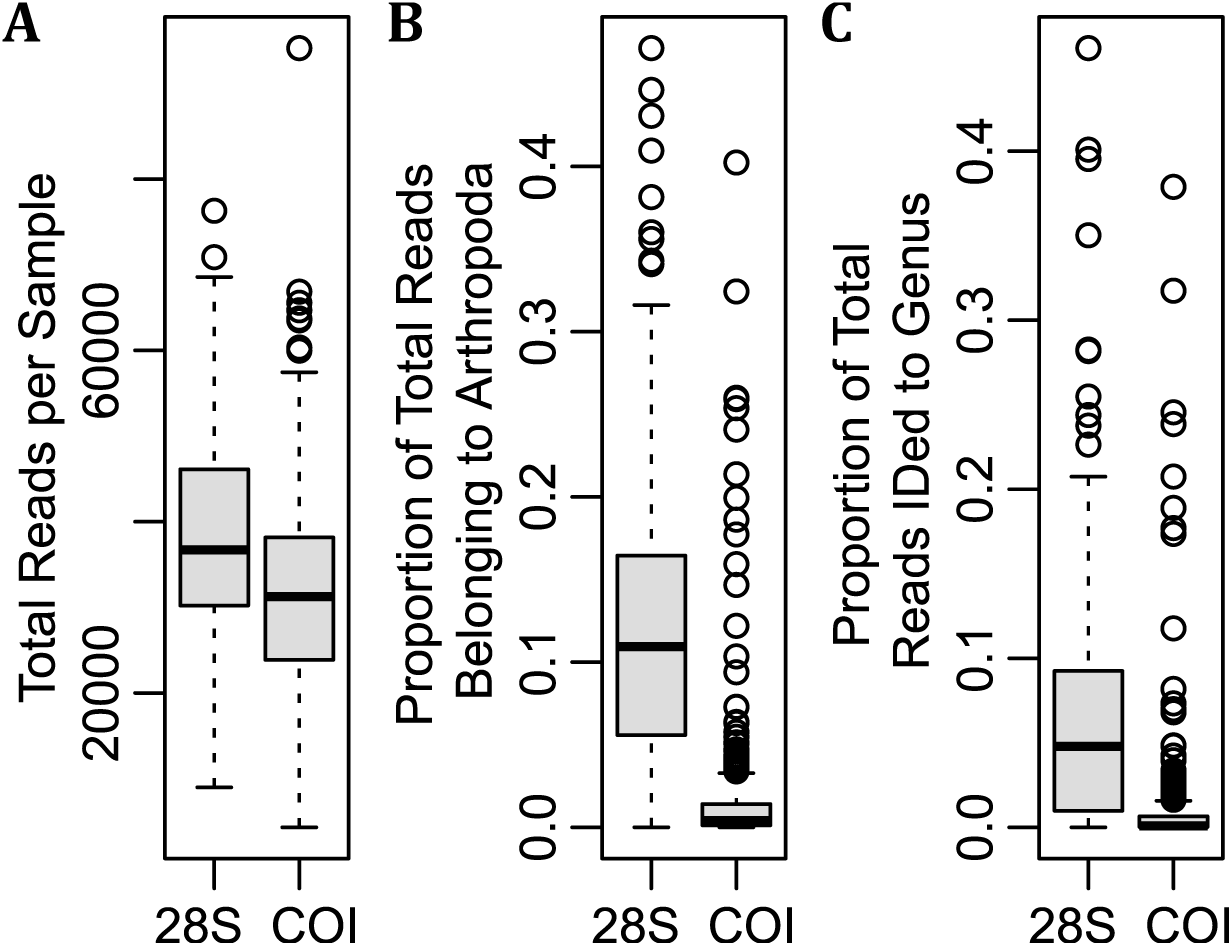
Analysis of per sample sequencing depths achieved (A), proportion of sequences identified as Arthropoda (B) and proportion of sequences identified to an arthropod genus (C) for both markers processed in this study.

In total, eDNA analysis of the collected samples produced 2,228 Arthropoda detections spanning 543 genera (Figure 3A). A large proportion of detections were represented by the orders Hymenoptera, Lepidoptera and Coleoptera. Hemiptera, Thysanoptera, Diptera and Araneae detections also featured prominently in the data. Similar patterns were observed regarding genus-level diversity of detections (Figure 3B), and prevalence of detection across samples (Figure 3C). Regarding Lepidoptera and Anthophila, groups of heightened conservation concern (Cornelisse et al., 2025), eDNA yielded a rich dataset with 558 Lepidoptera detections (21 families spanning 209 genera) 260 bee detections (5 families spanning 32 genera).

**Figure 3:**
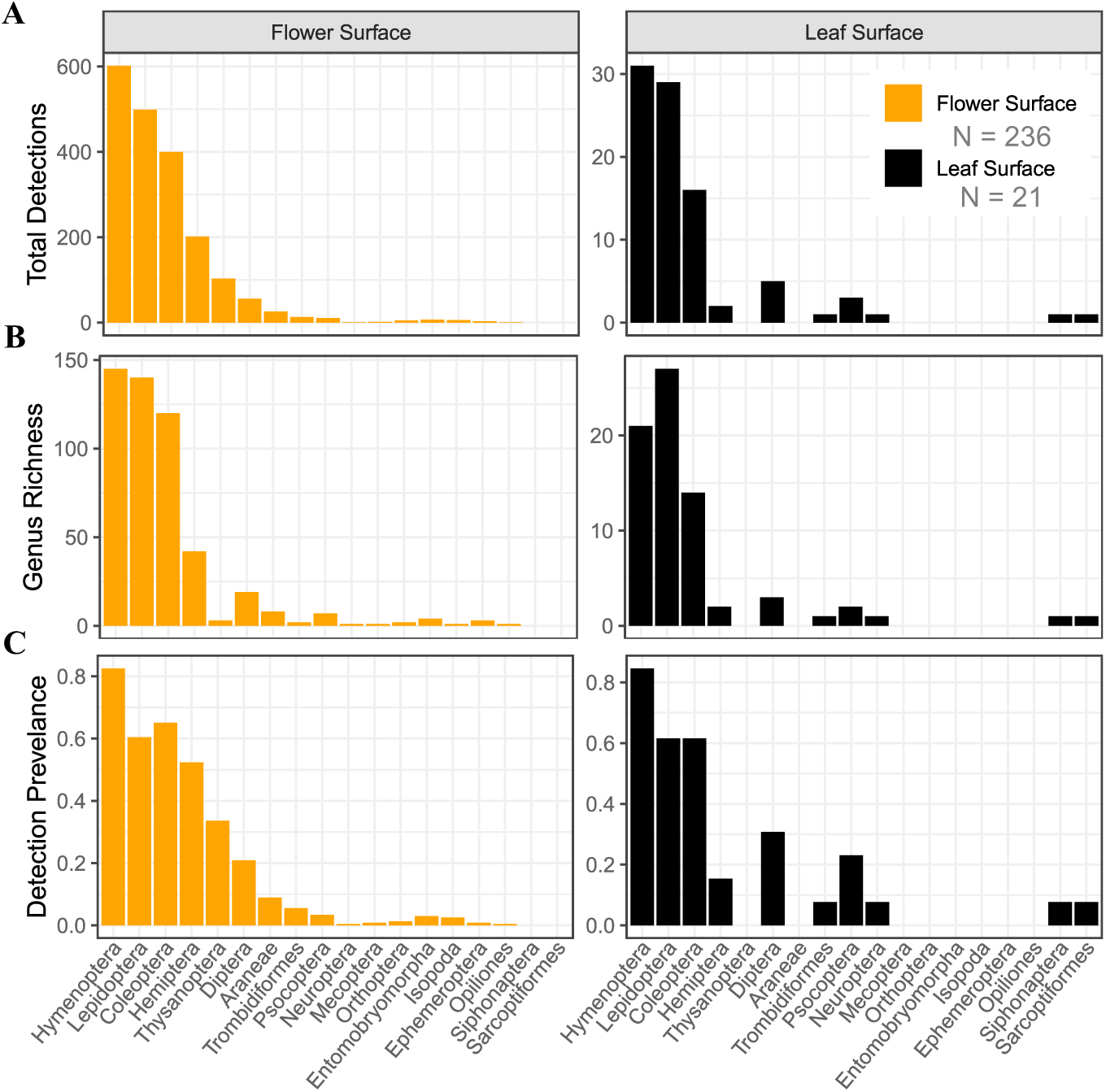
Summaries of study-wide genus-level eDNA detections (A), genus-level richness (B) and detection prevalence (C) across all arthropod orders detected, partitioned by flower surface and leaf surface samples.

In comparing trends across flower surface and leaf surface eDNA, we found no significant differences in the frequency of detection (*P* = 0.840) or genus-level richness per sample (*P* = 0.900) when analyzing all Arthropoda detections (Figure 4A). Similarly, we found no effects of substrate type when comparing frequency of detection (*P* = 0.754) or genus-level richness per sample (*P* = 0.489) when evaluating Lepidoptera (Figure 4B). However, we found an increased frequency of detection (*P* = 0.028) and increased genus richness per sample (*P* = 0.009) when evaluating results for Anthophila (Figure 4C).

**Figure 4:**
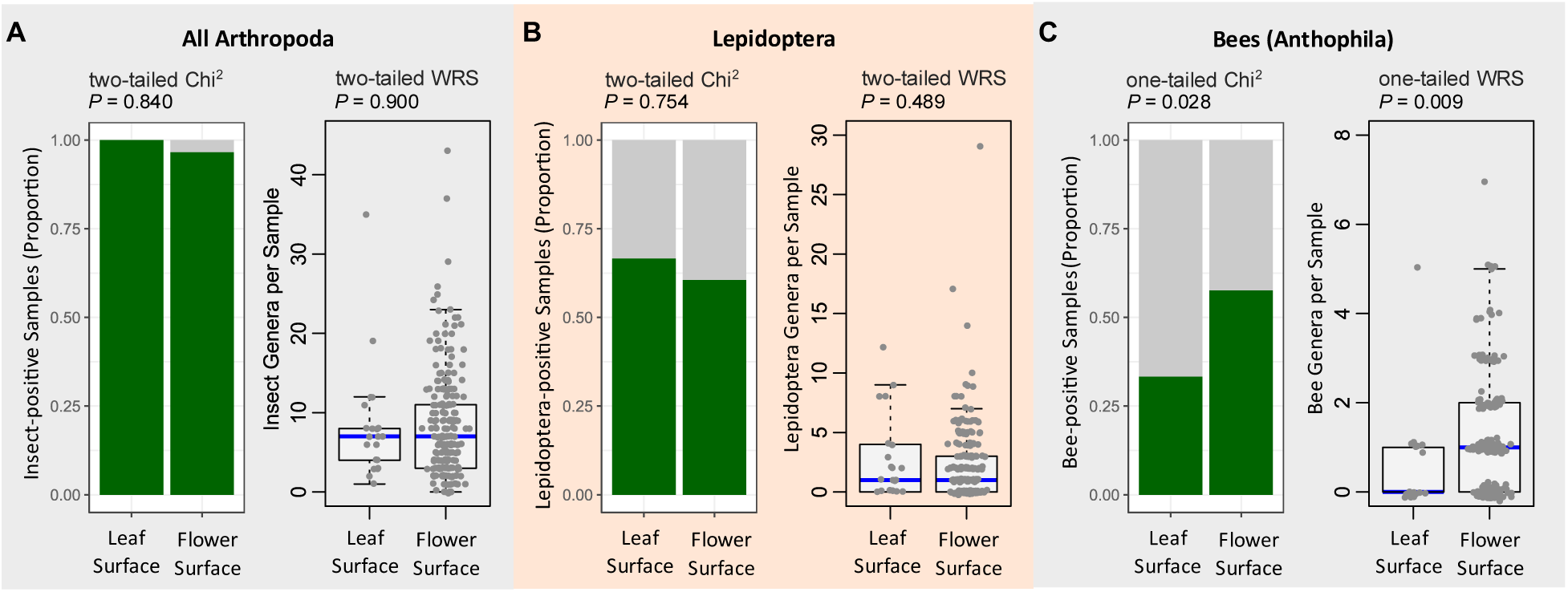
Comparison of flower and field negative eDNA samples. No statistical differences were observed in terms of frequency of detection of genus-level detection richness when conducting analysis for all Arthropoda (A) or all Lepidoptera (B). However, flower eDNA samples exhibited a significantly greater proportion of Anthophila (bee)-positive samples and greater richness of bee genera per sample relative to field negative controls (C).

During analysis of the associations between total arthropod richness and landscape-scale measures of forest habitat, initial model selection using a 3-km radius scale for forest habitat covariates resulted in the removal of *EvergreenForest* and yielded the model shown in Equation 2, where all predictors were found to be significant at an α of 0.05. When assessing the optimal landscape scale for modelling arthropod richness, the 3-km radius estimates were more strongly supported relative a 0.5-km (ΔWAIC = −14.50) or 1-km (ΔWAIC = −10.19) radius. There was significant evidence in support of all four predictors within this final model: *W* (*β* = −0.040, *P* = 0.012)*, D* (*β* = 0.261, *P* < 4.10e-12), *DeciduousForest* (Figure 5A, *β* = 0.857, *P* = 0.007), *ForestDiversity* (Figure 5B, *β* = 0.754, *P* = 0.018). Additional evaluation of this model revealed strong predictive power (Nakagawa and Schielzeth’s Conditional pseudo-*R^2^* = 0.721; Nakagawa & Schielzeth, 2013) and no evidence of variance inflation due to multi-collinearity (VIF < 1.05 for all predictors).

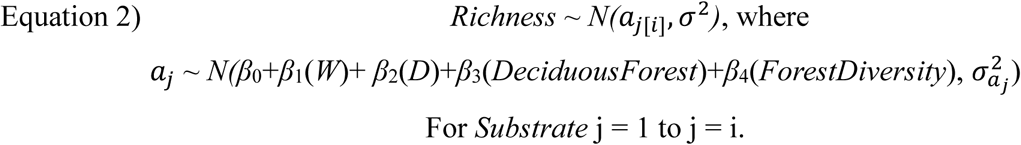

**Figure 5:**
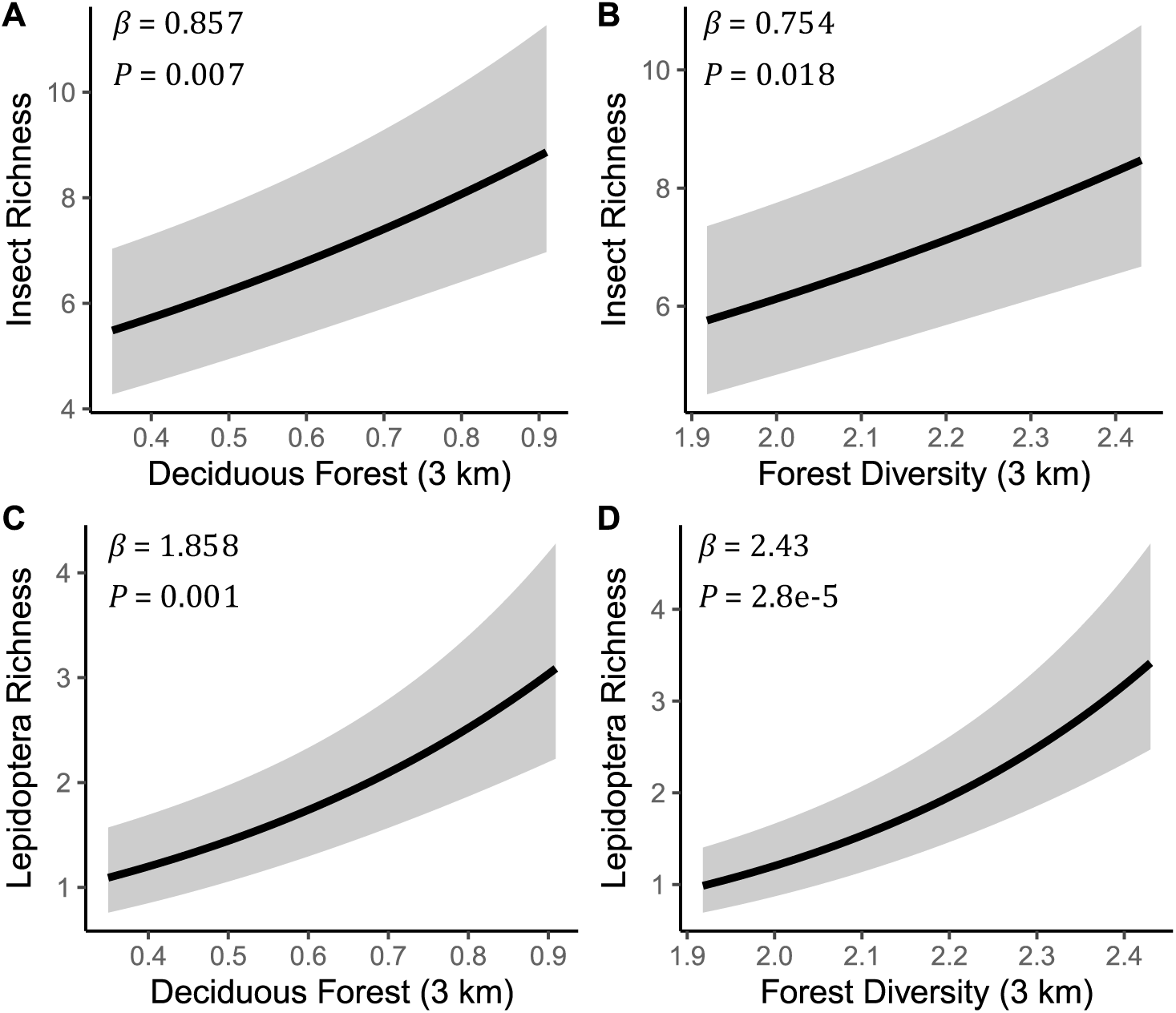
Partial effect plots of the best-fit relationships between eDNA-inferred sample richness values and measures of forest extent and diversity. Total insect richness was found to be significantly associated with the proportion of deciduous forest (A) and forest Shannon diversity (B). Among individual insect orders, Lepidoptera richness exhibited similar, yet stronger, associations with the proportion of deciduous forest (C) and forest Shannon diversity (D). The final WAIC-selected model for both total arthropod richness and Lepidoptera richness values is shown in Equation 2.

At the order level, analysis of Lepidoptera richness values revealed some of the largest effect sizes in the data. Model selection at a 3-km radius scale resulted in the removal of *EvergreenForest*, yielding the model shown in Equation 2. When assessing the optimal landscape scale for modelling Lepidoptera richness values, the 3-km radius estimates were more strongly supported relative to the 0.5-km (ΔWAIC = −57.38) or 1-km (ΔWAIC = −31.33). In the final model, *W* (*β* = 0.055, *P* = 0.055), *D* (*β* = 0.359, *P* = 2.40e-7), *DeciduousForest* (Figure 5C, *β* = 1.858, *P* = 0.001) and *ForestDiversity* (Figure 5D, *β* = 2.43, *P* = 2.80e-5) were all well-supported predictors of Lepidoptera richness within eDNA samples and follow-up evaluations revealed strong predictive power (Conditional pseudo-*R^2^* = 0.550) and no evidence of variance inflation (VIF < 1.05 for all predictors).

Modelling of Hymenoptera richness values resulted in the removal of *DeciduousForest* and *ForestDiversity*, yielding the model shown in Equation 3. Subsequent analysis supported a 0.5-km landscape scale relative to 1-km (ΔWAIC = −1.43) or 3-km (ΔWAIC = −2.95) scale. There was strong support for all three terms, *W* (*β* = −0.060, *P* = 9.30e-4), *EvergreenForest* (*β* = −0.859, *P* = 3.06e-3) and *D* (*β* = 0.290, *P* = 1.50e-11), and no evidence of variance inflation (VIF < 1.04 for all predictors).

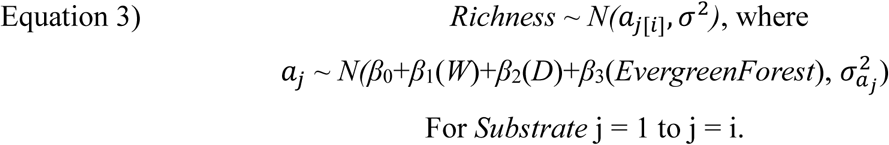

Given the strength of associations between forest habitat covariates and Lepidoptera richness, an additional round of model selection was conducted to evaluate whether insect richness values were related to forest habitat when Lepidoptera detections were excluded. As previously described, using Equation 1 as a starting model with forest covariates estimated at a 3-km radius, reverse step-wise selection resulted in the removal of *ForestDiversity* and *EvergreenForest*, yielding the model shown in Equation 4. Evaluation of scale revealed that a 0.5-km radius was more strongly supported relative to a 1-km (ΔWAIC = −7.93) or 3-km (ΔWAIC = −6.48) radius for the remaining forest habitat covariate. Within the final model, predictors *W* (*β* = −0.076, *P* = 7.10e-7), *D* (*β* = 0.224, *P* < 3.80e-10) and *DeciduousForest* (*β* = 0.474, *P* = 0.02) were all significant predictors of non-Lepidoptera arthropod richness.

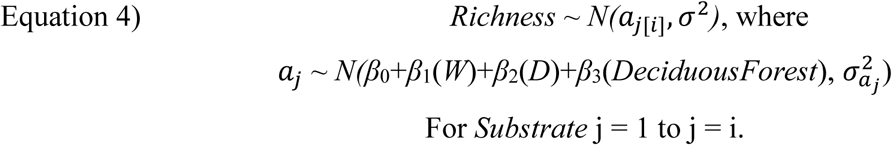

## Discussion

Our study developed novel insect eDNA methods for terrestrial systems, where biodiversity is difficult to assess at large scales. Using eDNA, we show that insect richness patterns were strongly predicted by surrounding landscape features including forest extent, both deciduous and mixed evergreen, as well as overall tree species diversity. These results provide a quantitative perspective on the current value of forest ecosystems for arthropod conservation within Central Appalachia. Our conclusions also highlight a potential mechanism through which past and future alterations to forested systems may influence arthropod conservation outcomes. This work will improve our capacity to track how anthropogenic change influences insect populations and guide insect decline mitigation efforts.

Implementation of eDNA methods resulted in high densities of detection and demonstrated clear capacity for characterizing arthropod-habitat associations. With that said, future efforts are warranted to continue improving eDNA applications. Consistent with past studies (Avalos et al., 2024; Jones et al., 2024), our findings highlight the powerful impact marker choice can have on breadth of detection. Even though there were approximately 2.5 times as many arthropod genera represented in the COI database relative to 28S, we detected 3 times as many arthropod genera using 28S relative to COI. This result is consistent with the observation that the proportion of 28S sequences belonging to Arthropoda was approximately an order of magnitude greater than that of COI. Ultimately, the BR2/BF2 COI primers showed minimal specificity for Arthropoda. Given the heightened power of the COI marker during species delimitation, development of COI primers with improved selectivity for Arthropoda would considerably improve methodological performance.

Relative to past arthropod eDNA studies, this work represents the largest sample size conducted to date (Table 1), reflecting the ease with which large numbers of samples can be collected and processed using the methods we describe. Unsurprisingly, this sampling effort resulted in more arthropod genera detected relative to past efforts. To produce a roughly standardized estimate of detection sensitivity, we divided the total arthropod genera detected in past arthropod eDNA studies by the square root of the sample size to account for the power-law relationship between sample size and species accrual (Macarthur & Wilson, 1967). In our data, this approach was strongly supported by a species accumulation analysis, where accumulated richness was linearly related to square root-transformed sample size (*R^2^* = 0.99, *P* < 0.001, OLS Regression). By this metric, our study, as well as two others (Allen et al., 2023; Stothut et al., 2024), stand out as highly sensitive implementations of arthropod eDNA surveys. Variation in performance across past works will likely guide future eDNA efforts towards greater levels of sensitivity. Of course, readers should note that these comparisons are only approximately standardized given differences in marker selection, bioinformatic processing and baseline sample diversity.

**Table 1:** Comparison of current peer-reviewed implementations of arthropod eDNA monitoring.

| Study | Sample size (N) | Mean sequencing depth | Fold difference in sequencing depth relative to this study | Total arthropod genera | Total genera / sqrt(N) |
| --- | --- | --- | --- | --- | --- |
| This study, both markers | 257 | 70,608 | 1.0X | 543 | 33.87 |
| This study, 28S only | 257 | 38,280 | 0.5X | 419 | 26.14 |
| Allen et al. (2023) | 40 | 174,911 | 2.5X | 353 | 55.81 |
| Stothut et al. (2024) | 31<sup>a</sup> | 32,073 | 0.5X | 156 | 28.02 |
| Thomsen & Sigsgaard (2019) | 57 | 5,029,448<sup>b</sup> | 71.2X | 113 | 14.97 |
| Avalos et al. (2024) | 27 | 73,447 | 1.0X | 58 | 11.16 |
| Johnson et al. (2023) | 37<sup>a</sup> | 2,779,670 | 39.4X | 68 | 11.18 |
| Newton et al. (2023) | 30<sup>a</sup> | 35,814<sup>b</sup> | 0.5X | 47 | 8.58 |
| Kestel et al. (2023) | 80 | 192,080 | 2.7X | 33 | 3.69 |
| Gamonal Gomez et al. (2023) | 60 | 22,049 | 0.3X | 10 | 1.29 |

<sup>a</sup> Some samples failed to amplify.
<sup>b</sup> Reported sequencing depth does not include sequences which were removed during quality filtering and is likely approximately 50 percent of actual sequencing depth.

Moving beyond coarse landscape categorizations, eDNA methods show promise for revealing the impacts of a broad range of environmental changes in global forest ecosystems. In the last century, eastern U.S. forests have undergone substantial changes that are expected to alter insect habitat suitability. Fire suppression has contributed to mesophication, driving turnover from oak and hickory forests to maple and beech-dominated stands (Alexander et al., 2021; Nowacki & Abrams, 2008). Invasive species introductions have resulted in the functional eradication of previously abundant species, including American chestnut *(Castanea dentata*; Hepting, 1974) and most species of ash (*Fraxinus* spp.; Herms & McCullough, 2014). Previous assessments suggest these trends have negatively impacted insect diversity (Sierzega & Eichholz, 2019; Wagner & Todd, 2016), yet our data indicate that Central Appalachian forests continue to exhibit heighted insect conservation potential. The Central Appalachian region remains one of the most intact, high-diversity forest systems in eastern U.S., with greater fire frequency and lower levels of mesophication relative to the broader region (Lafon et al., 2005; Nowacki & Abrams, 2008) and follow-up studies comparing these forests to those of the eastern U.S. more broadly are warranted.

Resolution of disagreements about the magnitude and complexity of insect decline requires monitoring to be conducted at large scales. We argue that eDNA methods could be central to achieving this goal. Large-scale studies would ideally be conducted using probability sampling techniques (Deming, 1975; Firth & Bennett, 1998) within an occupancy modelling framework to account for the potential of systematic bias in sampling location or detectability (MacKenzie et al., 2006). The cost-efficient and easily democratized nature of our methods increases the feasibility of scaling insect monitoring to meet these demands. Further validation of the accuracy and limits of eDNA methods against alternate sampling protocols is warranted, and we expect to see continued improvements to the sensitivity and comprehensiveness of future eDNA implementations.

## Acknowledgements

Efforts were primarily supported by a DoD ESTCP Grant to RTR and KG (Project RC22-B5-7373). Additional support was provided by Western EcoSystems Technology, Inc. and Metamorphic Ecological Research and Consulting, LLC. We thank USFWS, USFS, VA DCR-DNH, MD Forest Service, MD Park Service, MD DNR, WV DNR, WV State Parks and PA Game Commission. The authors thank Katherine Turo and John Lloyd for helpful manuscript reviews. Any use of trade, firm, or product names is for descriptive purposes only and does not imply endorsement by the U.S. Government.

## Supporting Data

All sequence data, reference sequence databases, and relevant summary tables needed to reproduce statistical analyses are publicly available at https://doi.org/10.5281/zenodo.15979887.

---

**Supplemental Table S1:**
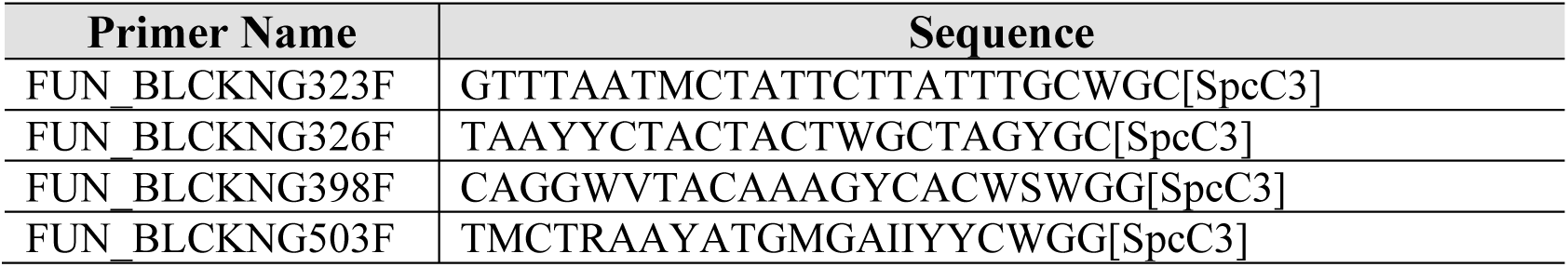
Blocking primers designed to suppress non-target fungal amplification.

